# Post-weaning social isolation alters sociability in a sex-specific manner

**DOI:** 10.1101/2024.07.11.603129

**Authors:** Teneisha Myers, Elizabeth A. Birmingham, Brigham T. Rhoads, Anna G. McGrath, Nylah A. Miles, Carmen B. Schuldt, Lisa A. Briand

**Author notes:** Equal author contribution (Teneisha Myers, Elizabeth A. Birmingham). **Address correspondence to:** Lisa A. Briand, Ph.D. Temple University Department of Psychology Weiss Hall, 1701 North 13^208^ Street Philadelphia, PA 19122.

## Abstract

Adolescence is a critical period for brain development in humans and stress exposure during this time can have lasting effects on behavior and brain development. Social isolation and loneliness are particularly salient stressors that lead to detrimental mental health outcomes particularly in females, although most of the preclinical work on social isolation has been done in male animals. Our lab has developed a model of post-weaning adolescent social isolation that leads to increased drug reward sensitivity and altered neuronal structure in limbic brain regions. The current study utilized this model to determine the impact of adolescent social isolation on a three-chamber social interaction task both during adolescence and adulthood. We found that while post-weaning isolation does not alter social interaction during adolescence (PND45), it has sex-specific effects on social interaction in adulthood (PND60), potentiating social interaction in male mice and decreasing it in female mice. As early life stress can activate microglia leading to alterations in neuronal pruning, we next examined the impact of inhibiting microglial activation with daily minocycline administration during the first three weeks of social isolation on these changes in social interaction. During adolescence, minocycline dampened social interaction in male mice, while having no effect in females. In contrast, during adulthood, minocycline did not alter the impact of adolescent social isolation in males, with socially isolated males exhibiting higher levels of social interaction compared to their group housed counterparts. In females, adolescent minocycline treatment reversed the effect of social isolation leading to increased social interaction in the social isolation group, mimicking what is seen in naïve males. Taken together, adolescent social isolation leads to sex-specific effects on social interaction in adulthood and adolescent minocycline treatment alters the effects of social isolation in females, but not males.

## 1. Introduction

Since the COVID-19 pandemic, adolescents are experiencing social isolation in ways that they have not previously. The level of social isolation experienced during the height of the COVID-19 pandemic has resulted in higher rates of depression and anxiety among young adults [1]. Conversely, high quality interactions with peers during adolescence in humans promote resilience later in life [2]. Rodent models have also demonstrated that social isolation, whether chronic or brief, has long lasting effects on brain development and behavior into adulthood. In rats, prolonged social isolation results in increased aggression towards conspecifics [3]. These changes in behavior are reflected in changes to the hypothalamic-pituitary-adrenal (HPA) axis and subsequent release of stress related hormones such as corticosterone in rodents [4]. In mice, social isolation increases anxiety-like behavior and promotes ethanol seeking behavior [5,6]. Additionally, these changes are more prominent in females than in males, as females experience higher rates of anxiety-like behavior than males following social isolation [7].

Post-weaning social isolation has effects far beyond behavior. Stressful experiences lead to changes in synaptic plasticity in various regions of the brain, and the brain is especially vulnerable to these changes during postnatal development [8]. Recent research has focused on microglia and their critical role in these changes. Microglia are a crucial part of the brain’s immune system, and chronic stress leads to increased activation of this system. Microglial activation following chronic stress causes morphological changes to microglia in which they become hyper-ramified. Hyper-ramification of microglia following chronic psychological stress is associated with behaviors that are representative of anhedonia and anxiety [9]. Examining the effects of chronic stress, such as adolescent social isolation, is critical in understanding how the brain is permanently changed over time, and how these changes can be attenuated.

The anti-inflammatory drug, minocycline, has shown success in inhibiting microglial growth. Minocycline administration during a period of prolonged stress successfully reverses the effects of chronic stress exposure on microglial ramification [10]. Inhibition of microglia through the administration of minocycline is not only effective in reversing the effects of stress, it also increases active coping behaviors in rodents [11]. Minocycline also encourages active recovery from social withdrawal [12]. The current study aims to examine how minocycline administration impacts the effects of adolescent social isolation stress on social behavior both in adolescence and in adulthood.

## 2. Material and Methods

### 2.1 Subjects

Male and female c57BL/6J mice were bred in house. On PND 21, at weaning, mice were randomly assigned to either isolation or group housing (3–5 mice per cage). All mice were held in an animal care facility with temperature and humidity control and kept on a 12 h light/dark cycle beginning at 7:30 am. The animals received food and water ad libitum and were given a cotton nestlet for enrichment. All procedures were approved by the Temple University Animal Care and Use Committee.

### 2.2 Drugs

Minocycline was purchased from Millipore Sigma (Burlington, MA) and was dissolved in sterile 0.9% saline. A dose of 40 mg/kg was delivered daily via intraperitoneal injection from postnatal day 21 to 42.

### 2.3 Three-chambered social interaction test

All animals were tested twice in a three-chambered social interaction test, once in adolescence (PND45) and once in adulthood (PND60). The apparatus consisted of a rectangular plexiglass three-chambered box with a lid. Each chamber was 16in. x 8in. x 9in with a 2in. x 3in. (high) door leading from the center chamber to either right or left side. Both right and left sides of the chamber contained a wire cage measuring 3.5in. in diameter and 7.5in. tall for either a novel partner or novel object to be placed in. Novel partners were age and sex matched to the test animal and the partner and object used at PND60 were different than the partner and object used at PND45. After being brought from the animal facility, the animals were allowed to habituate for twenty minutes to the dimly lit room (93 lumens). In the first portion of the test, the animals were allowed to freely explore the apparatus for twenty minutes. Time spent interacting with the empty cylinders by sniffing was hand scored using AnyMaze tracking software. The animal was then briefly removed while the novel partner and novel object were placed in the cylinders. The side which contained the novel partner vs. novel object were randomized in a counter-balanced fashion. The animal was placed back into the middle chamber of the apparatus and allowed to freely explore for ten minutes. Time spent interacting with the cylinders by sniffing was again hand scored using AnyMaze tracking software.

### 2.4 Adolescent Drug Treatment

A subset of animals received daily intraperitoneal (I.P.) injections of minocycline or saline for 21 days starting at PND21. Mice were weighed daily at the time of injection to insure proper dosing. We did not observe a significant difference in social interaction between the unhandled animals compared to the saline animals [exploration time during habituation, t(75)= 0.015, p=.98; exploration during social phase, t(75)=0.72, p=.47] therefore, the data from the unhandled and saline animals were combined.

### 2.5 Data Analysis

Analyses were performed using GraphPad Prism 9.5 Software. All data from the social interaction experiments were analyzed using repeated measures two-way ANOVAs with housing condition and test phase as the independent variables and social preference score as the dependent variable. The social preference score was calculated by subtracting the time the experimental animal spent sniffing the novel object cylinder from the time the experimental animal spent sniffing the novel partner cylinder. For the habituation phase both cylinders are empty but the preference score is based on the preference for the cylinder that would contain the novel partner during the social phase of the test. Sidak’s post hoc comparisons were made when main effects or interactions were detected (p < 0.05).

## 3. Results

### 3.1 Post weaning social isolation does not alter sociability in adolescence

Group housed (GH) and socially isolated (SI) male mice exhibit a similar social preference at postnatal day 45 (PND45) [effect of test: *F*(1,38)=23.2, *p*<.0001; effect of housing: *F*(1,38)=0.774, *p*=0.39; interaction: *F*(1,38)=0.547, *p*=0.46; Figure 1b]. Similarly, at PND45, female mice exhibit a social preference regardless of whether they were group housed or socially isolated at weaning [effect of test: *F*(1,35)=35.6, *p*<.0001; effect of housing: *F*(1,35)=0.0359, *p*=0.85; Interaction: *F*(1,35)=1.77, *p* = 0.19; Figure 1c].

**Figure 1:**
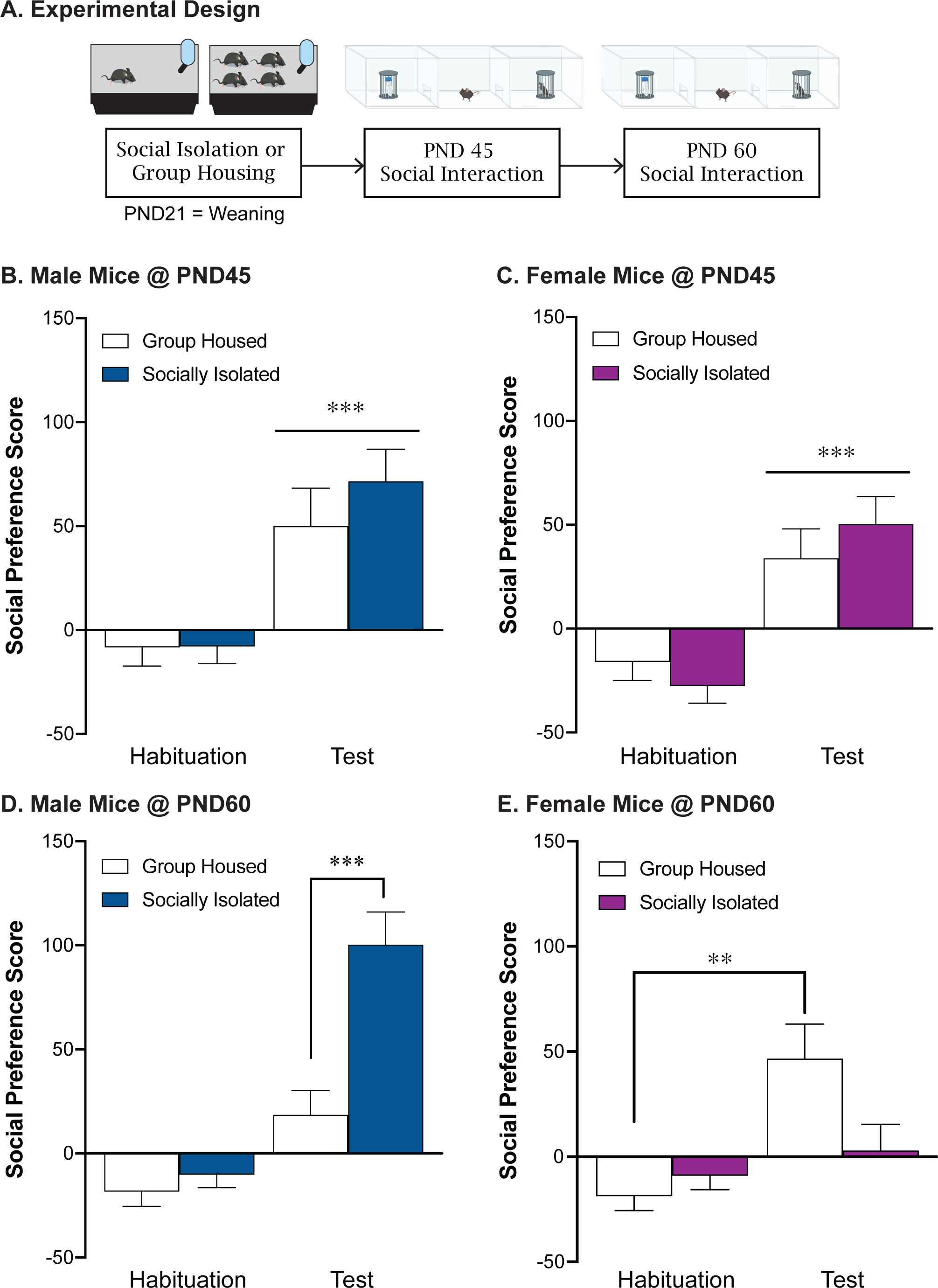
Post-weaning social isolation stress leads to sex-specific effects on social behavior during adulthood. **A.** Experimental timeline. Post-weaning social isolation stress does not impact male (**B**) or female (**C**) social preference score at PND45. Post-weaning social isolation leads to a significant increase in social preference score in males (**D**) and a significant decrease in social preference score in females (**E**) at PND60. **p<.01, ***p<.001

### 3.2 Post weaning social isolation has sex-specific effects on sociability in adulthood

During adulthood on postnatal day 60 (PND60), the socially isolated males displayed significantly greater social preference than group housed controls [effect of housing: *F*(1,38)=13.9, *p*=0.0006; effect of test: *F*(1,38)=60.13, *p*<0.0001; interaction: *F*(1,38)=14.9, *p*=0.0004; post-hoc test: GH test vs SI test: *p*<0.0001; Figure 1d]. In contrast, adolescent social isolation decreased social preference in females at PND60 compared to group housed controls [effect of housing: *F*(1,35)=7.12, *p*=0.178; effect of test: *F*(1,35)=46.1, *p*<0.0001; interaction: *F*(1,35)=4.29, p=0.046; post-hoc test: GH test vs SI test: *p*=0.001; Figure 1e]. When we statistically compare the social preference score during the test phase across males and females we see a significant interaction between biological sex and housing condition [effect of housing: *F*(1,73)=1.69, *p*=0.198; effect of sex: *F*(1,73)=5.65, *p*=0.02; interaction: *F*(1,73)=18.45, *p*<0.0001; post-hoc test: male GH vs male SI: *p*=0.001; female GH vs. female SI: *p*=.04; male SI vs. female SI: *p*<.0001].

### 3.3 Adolescent minocycline treatment decreases sociability in males but not females

Adolescent minocycline treatment dampened social preference in adolescent males at PND45 [effect of test: *F*(1,20)=4.21, *p*=.053; effect of housing: *F*(1,20)=0.007, *p*=0.93; interaction: *F*(1,20)=1.31, *p*=0.27; Figure 2b]. However, adolescent minocycline treatment did not alter social preference in females at PND45 [effect of test: *F*(1,19)=34.8, *p*<.0001; effect of housing: *F*(1,19)=1.07, *p*=0.31; interaction: *F*(1,19)=0.26, *p*=0.61; Figure 2c].

**Figure 2:**
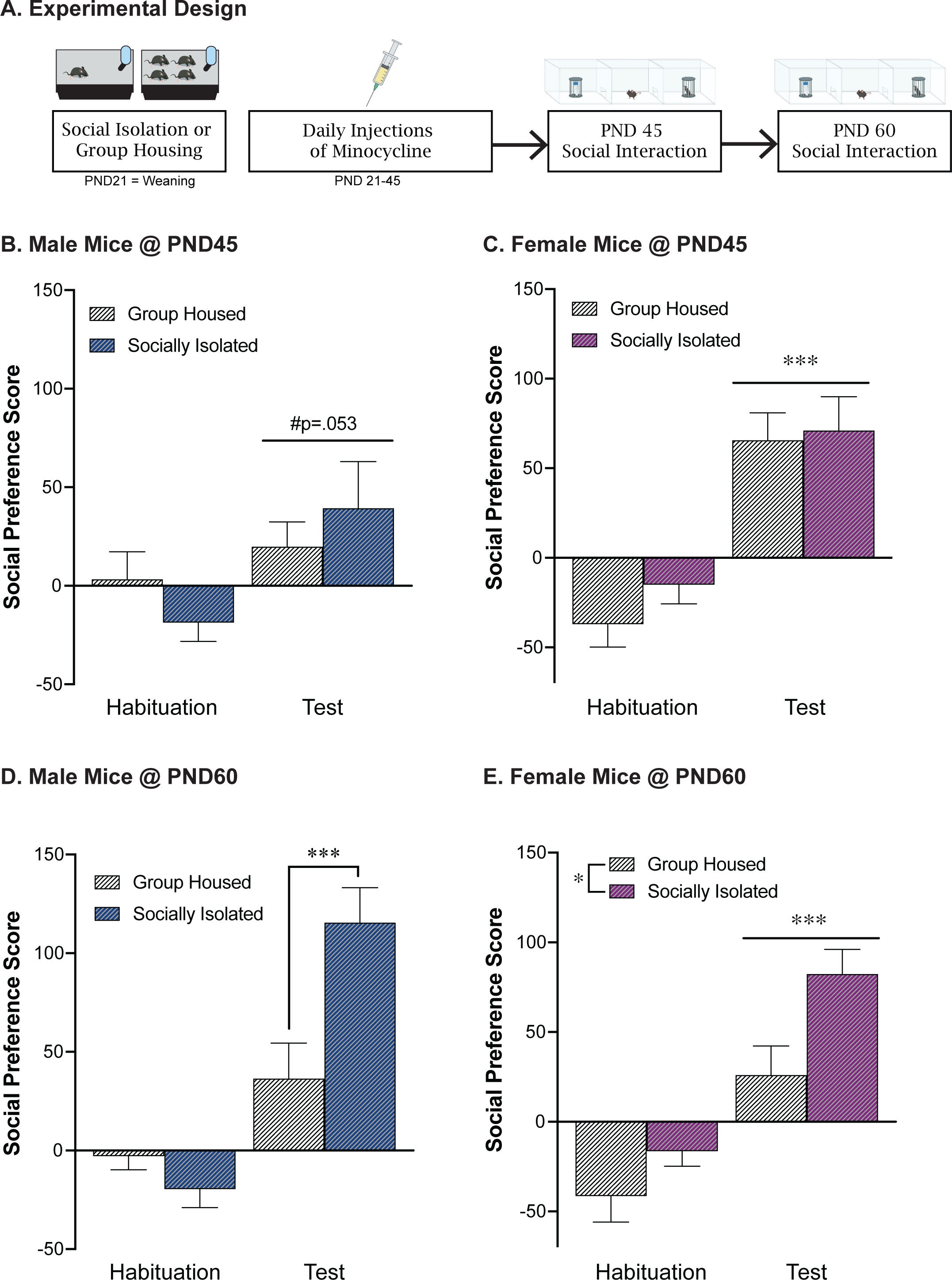
Adolescent minocycline treatment reverses the impact of post-weaning social isolation stress on social behavior in females during adulthood. **A.** Experimental timeline. Adolescent minocycline treatment decreases social preference score in males (**B**) at PND 45, while not impacting social preference in females at this timepoint (**C**). At PND60, adolescent minocycline treatment did not alter the impact of post-weaning social isolation, as isolated males continued to exhibit an increase in social preference (**D**). In contrast, adolescent minocycline treatment reversed the impact of adolescent social isolation in females, with isolated females exhibiting an increase in social preference at PND60 compared to group housed females (**E**). *p<.05, ***p<.001

### 3.4 Adolescent minocycline treatment reverses impact of social isolation on sociability in females in adulthood

Similar to what was seen in our unhandled and saline treated male mice, postweaning social isolation led to an increase in social preference in adult male mice compared to group housed controls [effect of housing: *F*(1,20)=4.56, *p*=0.045; effect of test: *F*(1,20)=41.9, *p*<0.0001; interaction: *F*(1,20)=12.63, *p*=0.002; post-hoc test: GH test vs SI test: p=0.0006; Figure 2d]. In contrast to what is seen in our unhandled and saline treated female mice, following adolescent minocycline treatment, both group-housed and socially isolated females exhibited a significant social preference [effect of housing: *F*(1,20)=7.19, *p*=0.015; effect of test: *F*(1,20)=46.1, *p*<0.0001; interaction: *F*(1,20)=1.64, *p*=0.22; Figure 2e]. In contrast to what we see in the absence of minocycline, following minocycline treatment, when we statistically compare the social preference score during the test phase across males and females we do not see a significant interaction between biological sex and housing condition [effect of housing: *F*(1,39)=16.39, *p*=0.0002; effect of sex: *F*(1,39)=1.70, *p*=0.20; interaction: *F*(1,39)=0.46, *p*=0.50].

## 4. Discussion

Overall, the present study demonstrates that post-weaning social isolation stress leads to a sex-specific effect on sociability during adulthood. In male mice, adolescent social isolation potentiated social preference in adulthood whereas in female mice social preference was disrupted in adulthood. During adolescence, no effects of housing were seen suggesting that the social isolation is impacting a developmental process that occurs between PND45 and PND60. Along with these sex-specific effects of post-weaning social isolation, we also found sex differences in the ability of minocycline to alter these social phenotypes. While minocycline disrupted social preference in adolescent males regardless of housing condition, it had no impact on the ability of adolescent social isolation to increase sociability in adult males. In contrast, minocycline reversed the direction of the effects of post-weaning social isolation on adult social behavior in females, with minocycline treated females exhibiting increased sociability following adolescent social isolation. This data suggests a sex-specific role of microglia that differs over developmental time periods.

### 4.1 Post-weaning social isolation stress leads to sex-specific effects on social behavior during adulthood

Adolescent social isolation leads to structural and functional changes in the brain [13–15]. Chronic stress during this developmental period further increases one’s susceptibility to neuropsychiatric diseases in adulthood, including anxiety, affective, and alcohol and substance use disorders [16]. Many stress-related psychiatric diseases are more prevalent in women [17,18] and understanding the potential sex-specific impacts of adolescent stress can provide models to further explore the brain mechanisms underlying stress-induced vulnerability.

The present study investigated the effects of post-weaning social isolation on social behavior in males and females. We found that post-weaning social isolation led to potentiated social preference in male mice in adulthood at PND60. Other groups have demonstrated increased social preference following post-weaning social isolation in males [16,19]. In addition, other models of post-weaning stress in male mice have also led to an increase in social preference in adulthood. Repeated footshock stress at PND21 led to an increase in social interaction in males in adulthood [20]. Although there is some congruence in the literature, not all studies examining the impact of post-weaning social isolation on social preference find consistent results [21]. These discrepancies may be impacted by the age at testing. In the current study we find no impact of social isolation at PND45 and studies that have examined social interaction at older time points [16,19] are more likely to see increases is social interaction compared to those that look earlier [21]. Additionally, many of these other studies have examined a battery of behavioral tests including tests that elicit anxiety-like behavior [16,21] and the timing of these tests relative to the social interaction testing could influence these results [22]. Differences could also be driven by differences in novelty preference as the social interaction procedure as the current study compared social interaction to exploration of a novel object rather than an empty cylinder or chamber as done in many other studies [16,20,21].

In contrast to what we saw in males, the current study found that post-weaning social isolation decreased social preference in adult female mice. One factor that may be mediating these effects is the impact of social hierarchy. While social dominance hierarchies alter the impact of social reward in males such that subordinate males find social interaction less rewarding, this is not the case in females [23]. Therefore, the differences in social interaction seen in the group housed animals may be mediated, in part, by social hierarchy and the lack of hierarchical experience may increase social reward in males.

Our work adds to the growing literature examining the role of isolation stress on the long-term behavioral impairments on social behavior in females [24–26]. The sex differences seen in the current study are consistent with the sex-specific outcomes of social stress in humans [7,17,18,27] and they point towards different neurological sequelae in response to social isolation. In fact, adolescence social isolation leads to sex-specific alterations in gene expression within the prefrontal cortex, nucleus accumbens and ventral tegmental area [28]. Overall, our data here suggests that social isolation during adolescence leads to opposing behavioral effects during adulthood in males and females and provides further support for sex differences in the mechanisms driving social stress induced behavioral alterations.

### 4.2 Post-weaning social isolation stress does not impact social behavior during adolescence

Given the literature on the sensitivity of the adolescent time period to chronic stressors impacting one’s behavior, we would expect to see post-weaning social isolation stress led to social behavioral impairments during this time. Interestingly, we did not observe this effect within the present study.

We found that males and females did not display a significant difference in their social preference when tested during adolescence compared to their group-housed controls. While this may seem contradictory to previous work, few studies exploring the effects of post-weaning social isolation stress have tested the animal’s social interaction behavior during the adolescent time point. Further this is consistent with work demonstrating increased “resilience” to stress during adolescence can manifest as alterations later in adulthood [29].

With these results seen in the present study, we expand upon the importance of examining social behavior at various timepoints to better the understanding of behavioral changes produced by isolation stress, as well as understanding the periods of development that are most sensitive to the stress exposure. Although adolescence has been established as critical period of development in its relation to stress sensitivity, the effects may show greater impact on long-term effects on brain development and behavior, with increasing prevalence during adulthood. With this, more research is needed investigating the impact of early social isolation stress on adolescent social behavior in both males and females [22,30,31]

### 4.3 Adolescent minocycline treatment reverses decrease in social preference in adult females following post-weaning social isolation

It is well established that early life stress leads to alterations of brain structure and function, which have the capacity to induce behavioral changes. Of the brain structures that are altered, microglia exhibit increased vulnerability to psychosocial stress exposure [9,10,32–36]. Social stress during adolescence leads to changes in microglial structure and function [37–41], with changes playing an important role in mediating social behavior [42,43]. Further, the impact of stress during adolescence on microglia is often sex-specific, with females exhibiting more stress-induced increases in microglial complexity than males [44]. Although highly complex, ramified microglia have long been considered quiescent, following chronic stress, microglia exhibit a hyper-ramified morphological state that is correlated with stress-induced behavioral alterations [43,45,46]. One such behavioral alterations seen following stress-induced microglial activation, is decreased sociability [47–50]. Administration of minocycline, an antibiotic with anti-inflammatory properties, attenuates the over-activation of microglia [42,51,52]. Following stress exposure, minocycline administration can reverse the microglial hyper-ramification and, in turn, reverse stress-induced behavioral impairments [11,42,51].

Consistent with this previous work, we showed that adolescent minocycline treatment during adulthood was able to reverse the effect that post-weaning social isolation stress had on social behavior in the females. This effect of minocycline seen here is consistent with the literature reflecting minocycline’s ability to attenuate behavioral deficits following exposure to a stressor [11,53]. Our work contributes to the growing research demonstrating minocycline’s role in alleviating behavioral deficits through the inhibition of hyper-ramified microglia. Overall, we conclude that our work expands upon this growing literature analyzing the role of microglia on social behavior following stress exposure.

### 4.4 Adolescent minocycline treatment diminishes social preference in males during adolescence while not impacting adulthood behavior

Here we found that both the adolescent group-housed and socially isolated males exposed to adolescent minocycline treatment displayed a decrease in social preference compared to their baseline controls. While this result may seem to conflict with previous findings demonstrating minocycline’s protective effects following behavior impairments, administration of the drug has shown to lead to adverse reactions in some cases, potentially attributed to antibiotic induced shifts in the microbiota [42,54,55]. In addition to the potential aversive effects, minocycline treatment has also shown to lead to reductions in locomotor behavior in rodents [56–58]. Taken together, this may explain why the adolescent males of the present study display a decrease in social preference, as the treatment may have led to the animals experiencing adverse reactions to the drug, leading to a decrease in locomotor behavior, further impacting their sociability.

### 4.5 Adolescent minocycline treatment leads to sex-specific alterations in social behavior

Taken together, we find that the effects of adolescent minocycline administration are sex specific. While the majority of preclinical studies that have examined the impact of stress on neuroimmune signaling have examined only male animals [9,59,60], human subjects research suggests that women may be particularly vulnerable to chronic stress exposure, including social isolation [7]. Preclinically, chronic stress often leads to sex specific alterations in neuroimmune signaling [44,61–63]. During development, microglial colonization differs across sex, with males exhibiting higher levels of microglial colonization early in life (PND4) and this flipping at PND30 to females exhibiting more microglia maintained in an activated state than males [64]. Therefore, female mice may be more vulnerable to manipulations the alter neuroimmune function, like social isolation, during this time period whereas male mice would be more vulnerable earlier. This is supported by work demonstrating that during early postnatal development, microglial phagocytosis plays a critical role in the development of social play behavior in males but not females [65,66]. The current study supports a role for microglia in adolescence in shaping adult female social behavior as inhibiting microglia with minocycline administration during adolescence blocked the ability of social isolation to disrupt social behavior, whereas the augmented social interaction seen in male mice following social interaction is not driven by microglial function.

## 4.1 Conclusions

Overall, the current study has shown that post-weaning social isolation stress leads to a sex-specific effect on sociability during adulthood, with the socially isolated males expressing an increase in social behavior while the females express a decrease. Furthermore, when exposed to adolescent minocycline treatment, we find that during adolescence, the male’s experience a decrease in their social behavior, yet this does not carry over into adulthood as they show no effect on social behavior at this timepoint. Although, the females express the opposite, showing no effect on their social behavior during adolescence yet an increase in social behavior during adulthood. Our results seen here further support previous work highlighting minocycline’s ability to attenuate behavioral deficits following stress exposure. This suggests that minocycline may be a potential treatment method for behavioral impairments following early life stress.

## Ethics approval and consent to participate

All procedures using experimental animals were approved by Temple’s Institutional Animal Care & Use Committee.

## Consent for publication

All authors read and approved the final manuscript for publication.

## Availability of data and material

The datasets used and/or analyzed during the current study are available from the corresponding author on request.

## Author Credit Statement

**Tenisha Myers**: Investigation, Validation, Formal Analysis, Writing – Original Draft, Visualization. **Elizabeth Birmingham**: Investigation, Formal Analysis, Writing - Reviewing & Editing, Visualization. **Brigham Rhoads:** Investigation, Project Administration. **Anna McGrath:** Conceptualization, Methodology, Investigation, Project Administration. **Nylah Miles:** Investigation. **Carmen Schuldt:** Investigation. **Lisa Briand:** Conceptualization, Methodology, Formal Analysis, Writing – Reviewing & Editing, Supervision, Project Administration, Funding acquisition.

